# MCAK/Kif2C centromeric activity level tunes K-fiber turnover through distinct pathways

**DOI:** 10.1101/2025.02.16.638494

**Authors:** Mike Wagenbach, Juan Jesus Vicente, Linda Wordeman

## Abstract

MCAK/Kif2C is a microtubule-depolymerizing kinesin implicated in the correction of chromosome attachment errors. When eliminated from kinetochores, cells exhibit delayed congression and a modest increase in chromosome missegregation. Curiously, MCAK/Kif2C overexpression (OE) promotes these same defects. Both depletion and excess levels of centromeric MCAK/Kif2C increase acetylated tubulin levels in the spindle, suggesting an increase in k-fiber stability. We conclude that this is the likely mechanism for the increase in chromosome segregation errors observed in both of these antagonistic conditions. Reduced MCAK/Kif2C increased the tubulin ratio on the two faces of the kinetochore, suggesting a greater likelihood of erroneous lateral MT interactions. In contrast, excess MCAK/Kif2C reduced the tubulin ratio at the kinetochore, stabilizing end-on MT interactions that increase the IKD and ultimately culminate in excessive stabilization of K-fiber microtubules. Both of these conditions promote chromosome segregation errors.

**Graphical abstract:** 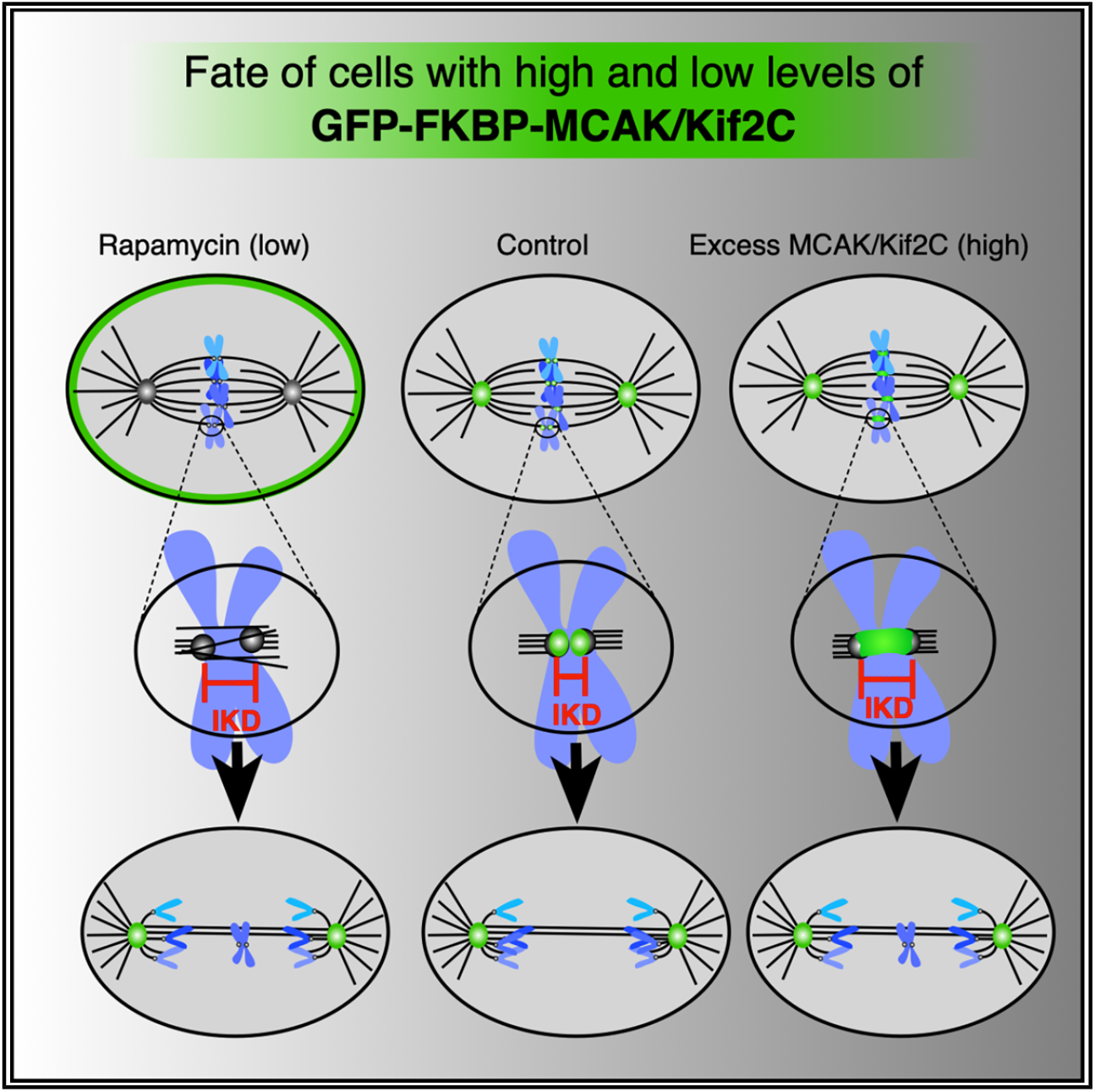

**In brief:** Cell biology; Chromosome organization

**Highlights:**

- An optimal level of MCAK/Kif2C ensures error-correction during mitosis
- Both excess or insufficient MCAK/Kif2C suppresses MT turnover in k-fibers
- Excess MCAK/Kif2C deforms kinetochores with uncoordinated movement
- Insufficient MCAK/Kif2C deforms kinetochores with unresolved lateral MTs

## INTRODUCTION

MCAK/Kif2C is the most well-studied member of the kinesin-13 family of microtubule (MT) depolymerizing kinesins. A comprehensive review of the protein’s enzymology, structure, and clinicopathology is available.^1^ Briefly, MCAK/Kif2C is thought to use its MT depolymerizing activity to detach aberrant MT connections from the kinetochore (KT), but it has never been determined, mechanistically, how these connections are recognized and severed by the enzyme. Thus, there is still much to be learned about this process. In cycling interphase cells, MCAK/ Kif2C is present in the cytoplasm, nucleus, and at the centrosome but disappears once cells differentiate terminally or become senescent.^2,3^ MCAK/Kif2C protein levels increase slowly up through G2, reaching peak levels at mitotic prophase and then returning to a minimal level in recently divided G1 cells.^4^ During mitosis, MCAK/Kif2C associates with a number of different cellular structures, including spindle poles, KTs, mid-bodies, and MT plus-ends.^5,6^ Studies have pinpointed a requirement for MCAK/Kif2C activity during congression to achieve metaphase chromosome alignment,^7–9^ although how this occurs is also unknown.

MT regulators such as MCAK/Kif2C present a challenge to study because when their levels are altered, tubulin homeostasis is perturbed, triggering compensatory changes in tubulin expression.^10^ We endeavored to circumvent this by changing MCAK/Kif2C levels (whenever possible) on a timescale that would not be permissive for compensatory cellular responses. To accomplish this, we have engineered CRISPR cells to express GFP-FKBP-MCAK/Kif2C under endogenous promoters. The protein can be relocalized within minutes in the presence of added rapamycin to any subcellular structure that is associated with exogenously added blue fluorescent protein (BFP) conjugated to the FKBP12-rapamycin binding (FRB) domain. By studying the enzyme at physiological levels in CRISPR-engineered cells, we have been able to add some refinements to our understanding of MCAK/Kif2C’s localization and behavior that differ from the published literature.

For example, in contrast to previous studies, including our own, MCAK/Kif2C is not found on MT plus-ends (+TIP) except during mitotic prometaphase. We do not see it on interphase MTs in cells expressing endogenous levels of MCAK/Kif2C.^11^ However, the protein can be artificially driven onto MT plus ends at any time during the cell cycle by expressing MCAK/ Kif2C from a strong promoter. Accordingly, we have found the quantity of +TIP-associated MCAK/Kif2C to be proportional to the cytoplasmic expression level of ectopic MCAK/Kif2C. Thus, any cells that have been transiently transfected with MCAK/Kif2C expressing constructs over and above the endogenous protein level will exhibit +TIP-associated MCAK/Kif2C when they normally would not. Another way to artificially drive MCAK/Kif2C onto MT plus-ends is by the ectopic expression of EB3, a protein not normally expressed in our cell lines. EB3 is preferentially expressed in muscle and neuronal tissue, but is often employed as a marker for MT plus ends when ectopically expressed in other cell lines.^12,13^ Interestingly, EB1, which is naturally expressed in our cell lines, does not artificially recruit MCAK/Kif2C to MT plus-ends even when overexpressed.^11^

We confirm that relocalization of MCAK/Kif2C results in an increase in lagging chromosomes at anaphase.^8,14,15^ However, because lagging chromosomes arise stochastically by, potentially, a variety of different mechanisms,^16–18^ this readout serves as a rather low-resolution method to determine the mechanistic contribution of MCAK/Kif2C to chromosome segregation. The limitation of the lagging chromosome assay is underscored by our observation of an increase in lagging chromosomes during both loss and overexpression (OE) of MCAK/Kif2C.^15^ One would think that these two antagonistic experimental treatments would surely lead to functionally distinct outcomes. The goal of this study is to mechanistically explain our observation that loss and OE exhibit the same phenotype. By employing the rapamycin-based ‘‘knock-sideways’’ approach to remove MCAK/Kif2C from mitotic structures within a timescale of minutes,^19^ we have been able to pinpoint with greater precision the time that MCAK activity is required during cell division: prometaphase. This is a time at which most, if not all, KTs are occupied by MCAK/Kif2C. In this study, we have used our CRISPR cell system to compare the mitotic effects of both MCAK/Kif2C loss and OE. While MCAK/Kif2C loss has been extensively investigated, OE has not.^15^ This is despite the fact that the protein is overexpressed in many cancers^20^ and, based on this study, has the potential to increase chromosome instability to promote tumor evolution.

Finally, in this study, we have used the ‘‘giant’’ X chromosome of the Indian muntjac cell line to develop means to predict, prior to anaphase, the likelihood that a chromosome possesses erroneous attachments. By normalizing our experimental regime to the X chromosome, we have measured the KT aspect ratio versus the tubulin ratio of the poleward and metaphase plate side of the KT to predict the type of aberrant connection that is associated with each experimental condition. We have confirmed that both loss and OE of MCAK/Kif2C delay chromosome congression to the metaphase plate to the same extent and concomitant with increased stability of KT fiber (K-fiber) MTs in our human cell lines. We believe that this occurs because we have found that increased inter-KT distances are promoted by both conditions. Increased IKD might place excess tension on the K-fiber MTs. Such pulling forces have been implicated in suppressing MT disassembly and turnover.^21,22^ We hypothesize here that the pulling forces that are evident in the absence of MCAK/Kif2C are supplied by lateral KT-MT interactions, whereas the pulling forces seen in MCAK/Kif2C OE are attributable to excess poleward force generation from end-on attachments.

## RESULTS

### Altered MCAK/Kif2C protein levels delay congression

We have engineered a cell line expressing GFP-FKBP-MCAK/ Kif2C with stably integrated BFP-FRB-SH (blue-fluorescent-protein-FRB-SH (membrane anchor)^19^ in the plasma membrane (Figure 1A, top). Briefly, the N-terminal sequence MGCIKSKG KDSA derived from the LYN kinase gene directs myristoylation of Gly-2 and palmitoylation of Cys-3 in the Golgi. These modifications ptomoyr accumulation of the acylated protein on the cytoplasmic face of the plasma membrane. We have treated this cell line in two different ways. By adding 1 µm rapamycin, we can rapidly relocalize GFP-FKBP-MCAK/Kif2C to the plasma membrane (Figure 1A, middle). Although we find complete removal of the membrane to occur within minutes, we apply rapamycin for 4 h prior to imaging for consistency between experiments. Sometimes, partial relocation occurs in certain cells that have partially or completely lost their FRB moiety. We do not include such cells in our analyses. Alternatively, we can transiently transfect a plasmid coding for either GFP-MCAK/- Kif2C or cherry-MCAK/Kif2C into cells expressing endogenous GFP-FKBP-MCAK/Kif2C (Figure 1A, bottom). When we use two GFP-linked constructs in this manner (usually to accommodate another fluorescent moiety such as our cherry-CENP-A expressing cells), we limit our analysis to cells with integrated fluorescence density that is twice that of an untransfected control. Alternatively, when possible, we can add excess Cherry-MCAK/- Kif2C on top of GFP-FKBP-MCAK/Kif2C (Figure S1A) up to 2-fold with fluorescence calibrated relative to Cherry-MCAK/Kif2C CRISPR-engineered cells. The GFP and cherry constructs are capable of heterodimerizing without any measurable changes in activity.^24^ We have not noticed any changes in localization between the two differentially labeled constructs. However, localization is dynamic during cell division and is not uniform between individual cells, making comparative quantification challenging. This prevents us from evaluating cells that are experiencing an extreme excess of MCAK/Kif2C while still enabling the reliable identification of transfected cells. We track and stage cells throughout cell division using fluorescently labeled DNA or a fluorescently CRISPR-engineered centromere marker CENP-A.

**Figure 1.**
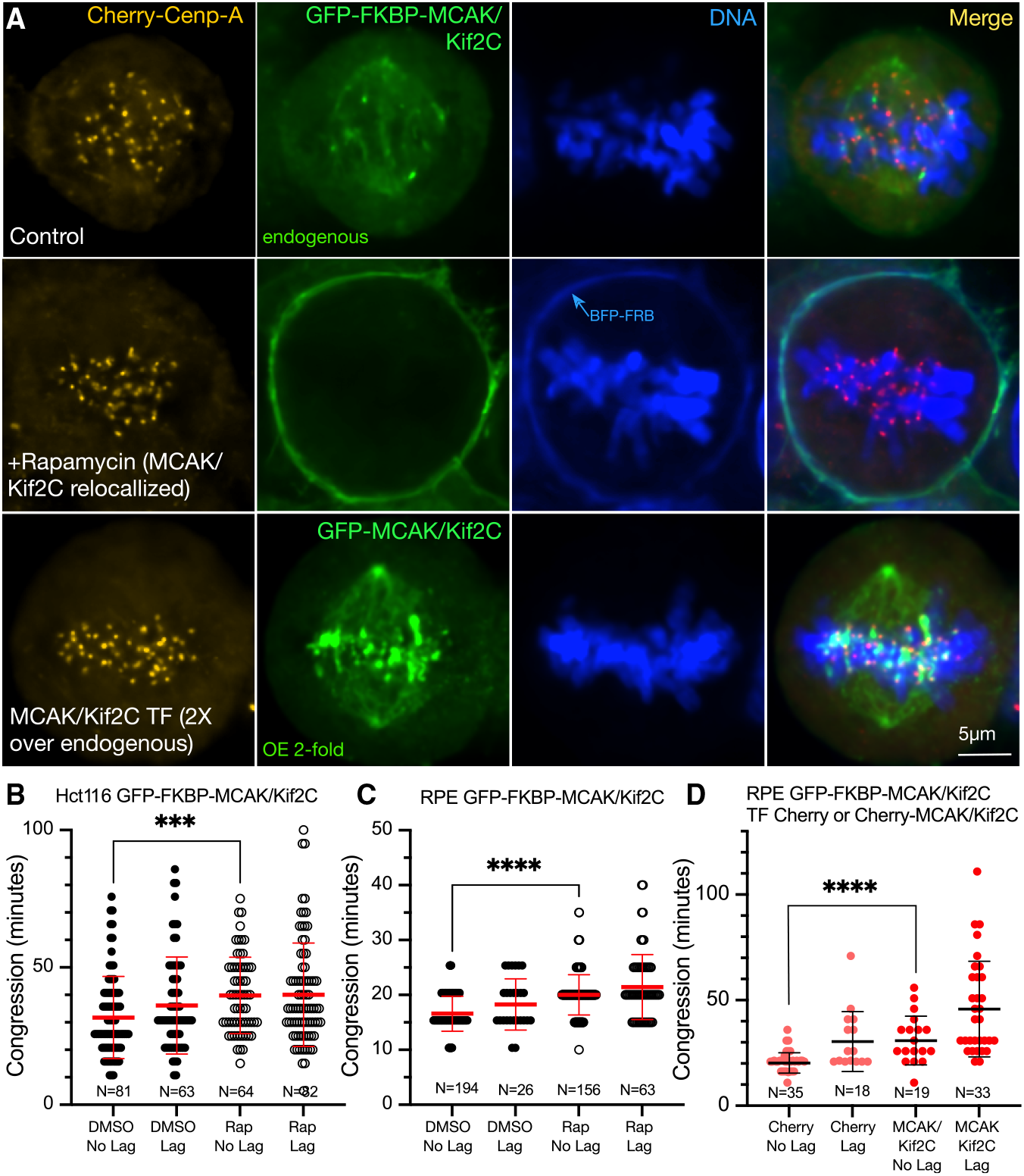
Live GFP-FKBP-MCAK/Kif2C cells exhibit delayed congression from NEB to metaphase when there is either too much or too little MCAK/Kif2C associated with the mitotic spindle. (A) GFP-FKBP-MCAK/Kif2C parent line co-engineered with cherry-CENP-A (top). GFP-FKBP-MCAK/Kif2C parent line stably expressing BFP-FRB-SH (middle), which relocalizes MCAK/ Kif2C to the plasma membrane in the presence of Rapamycin. An example of the GFP-FKBP-MCAK/Kif2C parent line transiently transfected with exogenous GFP-MCAK/Kif2C (bottom). The fluorescence level is 2-fold what would normally be seen in the parent cell line. (B) Live HCT116 GFP-FKBP-MCAK/Kif2C cells followed through mitosis in the presence or absence of rapamycin. \*\*\**p* = 0.0002; *N* = 81 cells from 4 technical replicates. (C) Live RPE GFP-FKBP-MCAK/Kif2C cells followed through mitosis in the presence or absence of rapamycin. \*\*\*\**p* < 0.0001; N = cells from 3 technical replicates. (D) Live RPE GFP-FKBP-MCAK/Kif2C transiently transfected with Cherry-MCAK/Kif2C. \*\*\*\**p* < 0. 0001; N = cells from 3 technical replicates. Cells in B, C, and D were followed through mitosis using Hoechst 33342. Scale bars, 5 microns for immunofluorescent images (A). All comparisons are Mann-Whitney t-tests.^23^

Live tracking of mitotic cells shows that both excess MCAK/ Kif2C and loss of MCAK/Kif2C delays the time required to progress from nuclear envelope breakdown (NEB) to the metaphase plate to a similar extent (Figure 1). On average, delayed congression is a regular feature of cells that go on to exhibit lagging chromosomes at anaphase (Figures 1B–1D, ‘‘Lag’’). However, we have found that congression is also delayed in cells that do not go on to exhibit lagging chromosomes but whose centromeres either lack MCAK/Kif2C (rapamycin-treated) (Figures 1B and 1C, closed vs. open circles) (*p* = 0.0002; *p* < 0.0001) or are associated with excess MCAK/Kif2C (transfected with either cherry-MCAK/ Kif2C (Figure 1D) (*p* < 0.0001) or GFP-MCAK/Kif2C (Figure 1A, bottom)). Statistical comparisons between parsed live cells that progress through cell division with no lagging chromosomes (Figures 1B–1D, Mann-Whitney *t* test, ‘‘no lag’’) are indicated.

Lagging chromosomes during anaphase are thought to primarily arise from unresolved merotelic attachments between one sister-KT and MTs emanating from both centrosomes. These connections arise naturally during the course of chromosome attachment to the mitotic spindle and congression.^25^ They are resolved prior to anaphase or even during anaphase in a mechanistically interesting but somewhat opaque manner.^26^

It is well-established from a number of studies (reviewed in^1^) that loss of MCAK/Kif2C can lead to a modest increase in lagging chromosomes during anaphase. This led to the idea that MCAK/ Kif2C is required to detach MTs from merotelic connections.

MCAK/Kif2C is an MT depolymerizer, so this activity would be expected to participate in the detachment of excess/erroneous MT attachments. However, it is puzzling that when MCAK/ Kif2C is only twice the endogenous levels (as measured by fluorescence intensity), cells present the same defect: lagging chromosomes and delayed congression.

MCAK/Kif2C associates with a number of mitotic structures (MT ends (during prometaphase), centrosomes, centromeres, KTs, and spindle MTs). Both the timing of centrosome separation and intrinsic MT assembly rates have been shown to adversely affect mitotic error correction.^27,28^ Accordingly, we tested whether centrosome separation (Figure S1B) or metaphase to anaphase timing when most MCAK/Kif2C has left the centromere (Figure S1C) was affected by MCAK/Kif2C loss and found no effect. Similarly, both interphase (Figure S2A) and mitotic MT assembly (Figure S2B) rates were not affected by MCAK/Kif2C loss. This led us to assume that the effect on congression and mitotic errors was primarily a function of centromere-associated MCAK/Kif2C.

### Anchoring the MCAK/Kif2C motor domain to the centromere can rescue congression defects

We tested this by anchoring either active (M) or inactive mutant (M^mut^) MCAK/Kif2C motor domain to centromeres using the centromere-binding domain of CENP-B. Note that this is only one of a number of regions where endogenous MCAK/Kif2C can localize within the centromere-KT domain.

The GFP-FKBP-MCAK/KIF2C parent cell line was transiently transfected with either Cherry-CPB-M^mut^ (serving as a control) or Cherry-CPB-M (possessing active MCAK/Kif2C activity). These constructs consist of 186–583 of the MCAK/Kif2C motor domain incorporating AAAAA mutations to prevent inactivation by Aurora B.^29^ Cherry-CPB-M^mut^ (pMX1400) incorporates the additional mutations H530A/R534A/K537A (‘‘HYPIR’’ mutations), which effectively inactivate the kinesin motor domain.^30^

When desired, cells transfected with CPB constructs can also be treated with rapamycin, which will relocalize the endogenous GFP-FKBP-MCAK/Kif2C to the plasma membrane while having no effect on the MCAK/Kif2C motor domain (M or M^mut^) anchored to the centromere. Representative fixed cells counterstained with Hoechst 33342 (Figures 2A and 2B; blue) and anti-centromere antibodies (Figures 2A and 2B; red) are shown. Congression times from live cells tracked through mitosis are shown in Figure 2C. The time between NEB and the establishment of the metaphase plate is plotted on the Y axis. Cells from each treatment were also parsed for the appearance of lagging chromosomes at anaphase. Notably, adding MCAK/Kif2C activity to centromeres in the presence of endogenous GFP-FKBP-MCAK/Kif2C delayed congression and produced chromosome segregation errors similar to what we see with OE of full-length MCAK/Kif2C (Figure 2C, CBP-M^mut^, salmon circles, vs. CBP-M, red circles, No Lag) (*p* = 0.0179). However, this condition is rescued if endogenous GFP-FKBP-MCAK/Kif2C is relocalized to the plasma membrane (Figure 2C, CBP-M^mut^, salmon circles, vs. CBP-M, No Lag, +Rap, red circles, green periphery) (ns, *p* = 0.362). A similar rescue is not seen when CBP-M^mut^, (a version of the MCAK/Kif2C motor domain lacking MT depolymerizing activity) is treated with rapamycin (*p* < 0.0001). Instead, the effect is identical to depleting MCAK/Kif2C in the parent cell line because the CPB-M^mut^ construct does not contribute MCAK/Kif2C activity to the centromere. Thus, centromere-localized MCAK/Kif2C is responsible for facilitating congression, and this function is disrupted either by loss or addition of excess centromere-associated MCAK/Kif2C activity above endogenous levels.

**Figure 2.**
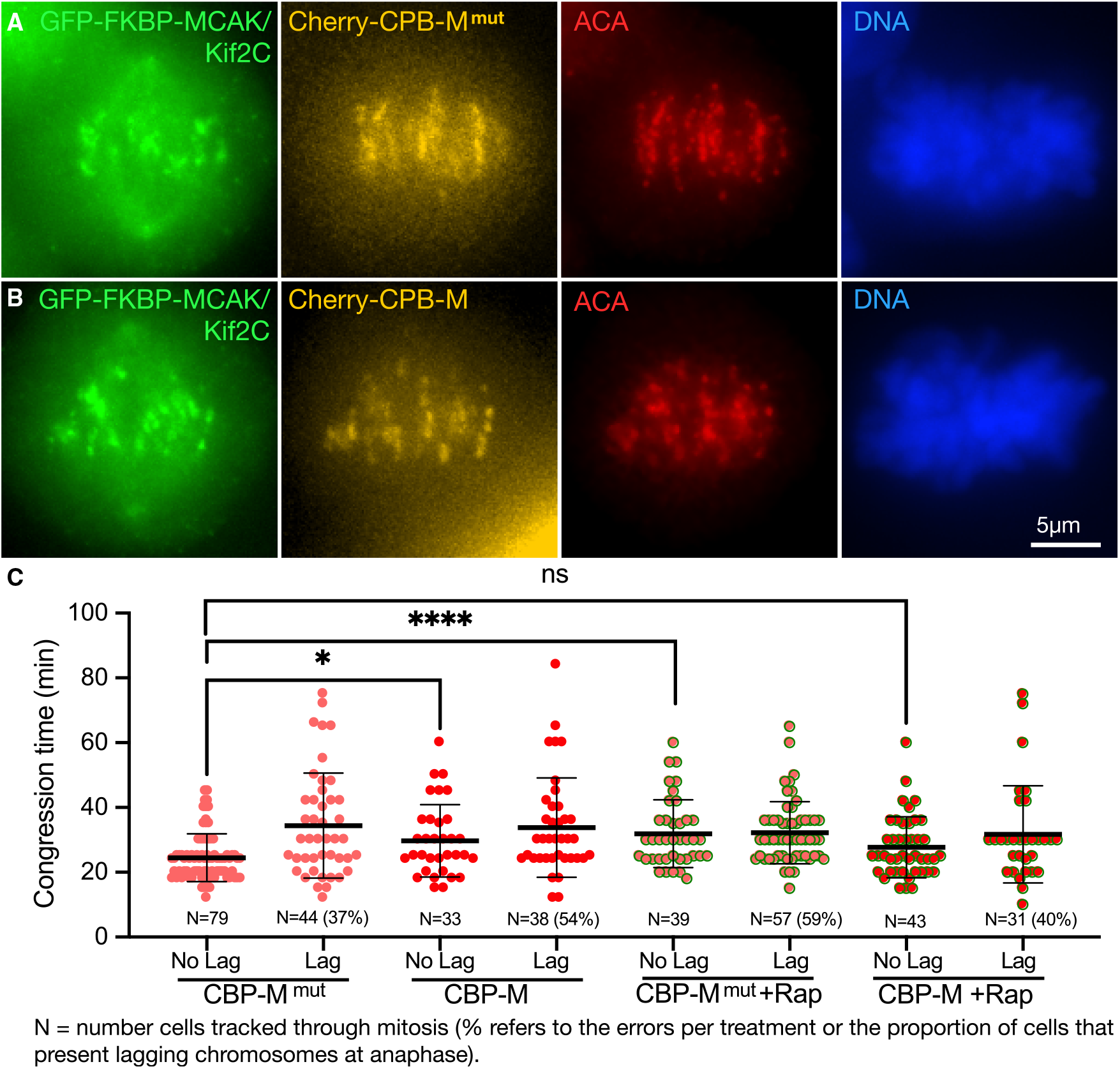
Anchoring the motor domain of MCAK/Kif2C to the centromere can rescue the congression and segregation defects associated with the loss of MCAK/Kif2C in rapamycin. (A) GFP-FKBP-MCAK/Kif2C parent line transfected with Cherry-CBP-M^mut^ (orange). Endogenous GFP-FKBP-MCAK/Kif2C (green). Immunofluorescently labeled centromeres (red) and DNA (blue) are shown. (B) GFP-FKBP-MCAK/Kif2C parent line transfected with Cherry-CBP-M (orange). M^mut^ = HYPIR mutated inactive motor (control), M = Active MCAK/Kif2C motor. (C) Live cells fluorescently tracked through mitosis. Congression times are plotted. Cells that exhibited lagging chromosomes are noted (Lag), and the percentage of tracked cells with lagging chromosomes is indicated (parentheses). ns, *p* = 0.362); \**p* = 0.0179; \*\*\*\**p* < 0.0001. N = individual cells from 3 to 5 technical replicates. All comparisons are Mann-Whitney t-tests.^23^

### Altered MCAK/Kif2C levels increase acetylated tubulin levels in the mitotic spindle

MCAK/Kif2C has been identified as important for congression by other researchers and we have confirmed and extended these data.^7–9^ However, the deleterious effect of excess centromere-associated MCAK/Kif2C has only been previously described by our laboratory.^15^ We are puzzled to find that BOTH loss and excess MCAK/Kif2C activity associated with the centromere delays congression and increases the percentage of lagging chromosomes. Because MCAK/Kif2C was thought to be associated with MT plus ends in interphase, we tested whether MCAK/Kif2C affected centromere separation but found no difference (Figure S1A). This is not surprising, as our CRISPR cells revealed that interphase plus tip localization is an artifact of EB3 coexpression.^11^ We have also tested metaphase-to-anaphase timing (Figure S1B) and have found this parameter to be unaffected by loss of centromeric MCAK/Kif2C. We did not further pursue centrosome separation or metaphase-to-anaphase timing. Instead, we focused on determining the mechanism by which these two conditions (loss and excess of centromeric MCAK/Kif2C) promote chromosome attachment errors and delay congression. To parse the deleterious characteristics of loss and OE of MCAK/Kif2C, we evaluated putative K-fiber stability using acetylated tubulin as a readout. We found that MCAK/Kif2C loss, excess MCAK/Kif2C, and excess centromere-associated active MCAK/Kif2C motor domain were each correlated with increased acetylated tubulin in the spindle (Figure 3).

**Figure 3.**
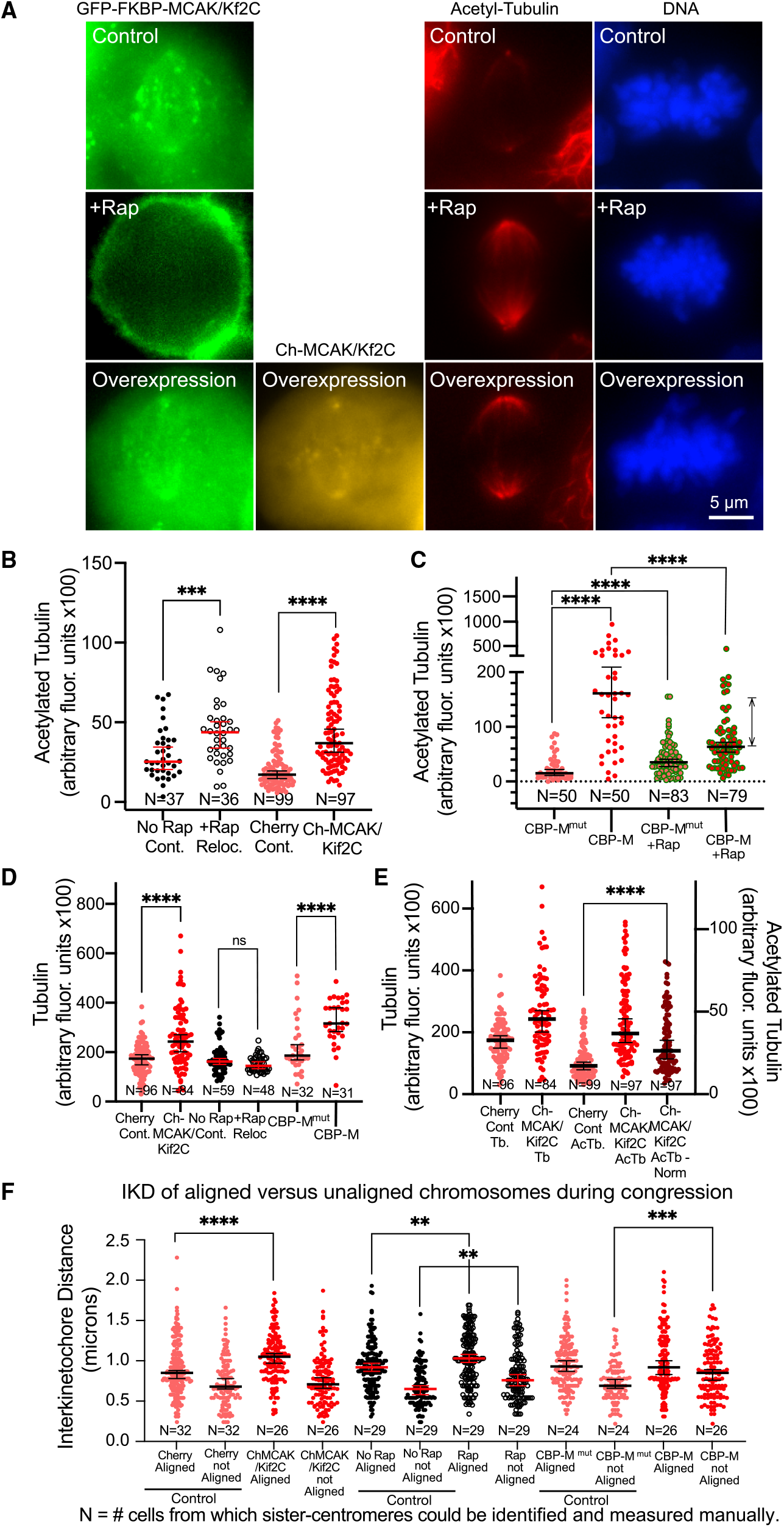
Both excess MCAK/Kif2C activity and loss of MCAK/Kif2C activity raise the level of acetylated tubulin in the spindle. (A) Representative fixed cells labeled with acetyl-tubulin in the far-red channel (second from right, red). GFP-FKBP-MCAK/Kif2C is shown (left), overexpressed cherry-MCAK/Kif2C (second from left, orange), and DNA (right, blue) are shown. (B) Loss (open circles \*\*\**p* = 0.0007) or excess (red circles \*\*\*\**p* < 0.0001) MCAK/Kif2C leads to higher levels of acetyl-tubulin in the spindle. (C) Anchoring excess MCAK/Kif2C activity to centromeres (red circles) significantly raises the level of acetyl-tubulin in the spindle. This can be partially rescued by removing endogenous MCAK/Kif2C from centromeres (red with green perimeter circles) \*\*\*\**p* < 0.0001. (D) Treatments that require overnight expression (Ch-MCAK, red, left, and CBP-M, red, right) also increase the level of tubulin in the spindle \*\*\*\**p* < 0.0001. Treatment with rapamycin (open circles) does not increase tubulin in the spindle. (E) Normalization of acetyl-tubulin expression relative to the increase in tubulin expression (red circles, left) does not eliminate the significance of the increase in acetyl-tubulin (red circles, right) caused by overnight MCAK/Kif2C expression (dark red circles, right) \*\*\*\**p* < 0.0001. (F) IKD in aligned centromeres and those two microns away from the spindle midzone (unaligned) ****, \*\**p* = 0.0014, \*\**p* = 0.0028, and \*\*\**p* = 0.0014. All comparisons are Mann-Whitney t-tests.^23^

Figure 3A shows representative fixed cells measured in this assay. Endogenous GFP-FKBP-MCAK/Kif2C is shown (green), along with excess cherry-MCAK/Kif2C (orange), where used. Cells were labeled in far-red with anti-acetyl-tubulin (red) and DNA (Hoechst 33342, blue). We observed that cells in which MCAK/Kif2C was removed (Figure 3B, open circles) (*p* = 0.0007) or overexpressed (Figure 3B, red circles, right) (*p* < 0.0001) or excess active motor anchored to centromeres (Figure 3C, red circles, left) (*p* < 0.0001) exhibited higher putative K-fiber-associated acetylated tubulin fluorescence. Representative images of cells expressing CBP-M and CBPMmut (Figure 3C) and labeled with antibodies against acetylated tubulin are shown (Figure S2C). Moreover, removal of endogenous GFP-FKBP-MCAK/Kif2C with rapamycin partially rescued (decreased) the acetyl-tubulin fluorescence levels in the spindle (Figure 3C, red circles with green periphery, right) (*p* < 0.0001) in agreement with the rescue of congression time and errors seen in this regime (Figure 2C).

Given that changes in IKD can appear to alter K-fiber stability, and influence K-fiber MT dynamics,^21,22,31^ we investigated the interkinetochore distance (IKD) between sister-centromeres in aligned chromosomes and also in chromosomes that have not yet reached the metaphase plate in naturally congressing chromosomes on bipolar spindles (Figure 3F). As has been shown numerous times, aligned chromosomes exhibit a larger IKD than those that have not yet aligned at the metaphase plate (Figure 3F, far left, salmon circles). Cells with excess MCAK/ Kif2C (Figure 3F, left, red circles) have higher IKD in aligned chromosomes relative to cells expressing cherry protein (Figure 3F, far left, salmon circles) (p<0.0001). Cells with no MCAK/Kif2C on centromeres (Figure 3F, black open circles), because it is relocalized in rapamycin, also exhibit higher IKD on aligned (*p* = 0.0014) or unaligned (*p* = 0.0028) chromosomes relative to control cells (Figure 3F, black circles). Cells with active MCAK/Kif2C anchored to the centromere (Figure 3F, right, red circles) exhibit a higher IKD relative to control cells (Figure 3F, right, salmon circles) but with the marked exception that this is manifest only on chromosomes prior to alignment (*p* = 0.0014).

### Tubulin ratios at the kinetochore face reflect different pathways that limit sister kinetochore co-ordination

We turned to male Indian muntjac (*Muntiacus muntjak*) cells^32^ to investigate in greater detail the differences between MCAK/ Kif2C OE and depletion. Male Indian muntjac cells possess 7 chromosomes, one of which is compound and quite large (Figure 4A). We have not produced CRISPR cells for relocalizing MmMCAK/Kif2C, but we have prepared custom siRNA that depletes endogenous MmMCAK/Kif2C (Figures 4B and 4C). Using this siRNA and OE of CgMCAK/Kif2C we measured spindle acetylation and found that, as in other cell lines, both loss and OE of MCAK/Kif2C significantly increase acetylation (Figure 4D) (*p* = 0.0087; *p* < 0.0001 respectively). Similarly, both treatments led to increased IKD (Figure 4E) (*p* < 0.0001; *p* = 0.020).

**Figure 4.**
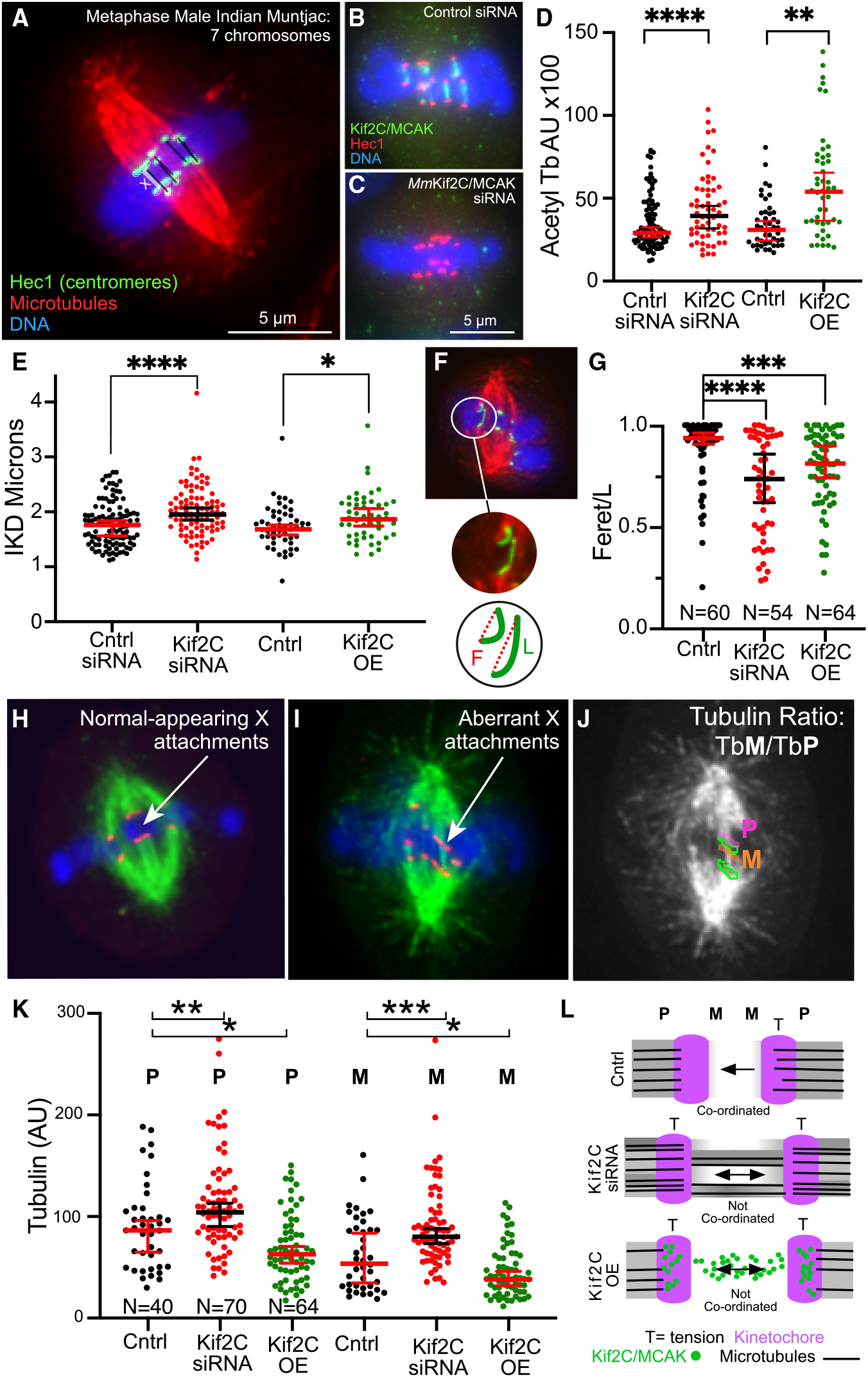
Tubulin ratios differ on each face of the kinetochore depending on the experimental treatment. (A) Representative fixed Muntiacus muntjak metaphase cell. Seven chromosomes are indicated, with the X chromosome indicated in white. Labels are indicated (lower left). (B) Mitotic Mmcell treated with control siRNA and labeled for MCAK/Kif2C (green). (C) Mmcell treated with siRNA directed against MmMCAK/Kif2C and labeled for MCAK/Kif2C (green). (D) Cntrl siRNA-treated cells (black circles, left), MCAK/Kif2C siRNA-treated cells (red circles), untransfected cells (black circles, right), and cells expressing excess GFP-MCAK/Kif2C (green circles) labeled for acetylated tubulin. \*\**p* = 0.0087; \*\*\*\**p* < 0.0001 respectively. (E) X chromosome IKD measurements in different experimental treatments. \*\*\*\**p* < 0.0001; \**p* = 0.020. Measurements are individual X chromosomes in 3 technical replicates. (F) Diagram of F/L measurements. (G) F/L is reduced in both loss (red circles) and excess MCAK/Kif2C (green circles). \*\*\*\**p* < 0.000 1; \*\*\**p* = 0.0009. (H) Metaphase Mmcell shows a normal appearing X chromosome centromere (red) attached to MTs (green). *N* = 54–64 individual X kinetochores in 3 separate replicates. (I) Abnormal appearing X chromosome centromere (red) attached to MTs (green). (J) Diagram of P (pink) and M (orange) tubulin measurements for one sister centromere (green). (K) P and M measurements for Cntrl (black circles, MCAK/Kif2C siRNA-treated (red circles), and MCAK/Kif2C overexpressing (green circles) X-centromere measurements. \**p* = 0.0052; \*\**p* = 0.0003; \*\*\**p* = 0.016; \**p* = 0.025. (L) Diagram showing how loss and OE of MCAK/Kif2C can lead to higher IKD and loss of coordination between sister centromeres. This condition could limit k-fiber turnover in both experimental conditions. All comparisons are Mann-Whitney t-tests.^23^

After confirming that this cell line behaves identically to our htert-RPE and HCT116 cells with respect to acetylation and IKD, we developed measurements that can be applied to the uniquely large X chromosome centromere to investigate attachment status. In the first, we measure Feret’s diameter (the longest straight distance between the two ends of a line traced along the centromere boundary, F) versus total centromere length (L) (Figure 4F). When F/L approaches ‘‘1’’ the centromere is straight and perpendicular to the long spindle axis. (see STAR Methods). After experimental treatment, we found that F/L is significantly reduced in both loss and increased MCAK/Kif2C (Figure 4G) (*p* < 0.0001; *p* = 0.0009). This indicates that the X centromere deviates from linearity under these conditions and constitutes a measure of centromere (and by extension, KT) deformation. We will refer to the centromere defined by the Hec1 label as the KT to distinguish it from the other regions of the K-fiber. We also measured the tubulin fluorescence (integrated density of identical areas) on the poleward (P) and metaphase plate facing (M) side of the X-KT (Figures 4H–4J). This is the sole measurement in which loss of MCAK/Kif2C differed from MCAK/Kif2C OE (Figure 4K). Both P and M tubulin measurements were increased in MCAK/Kif2C-depleted cells (*p* = 0.0052; *p* = 0.0003) (Figure 4K, red circles). In contrast, MCAK/ Kif2C OE reduced tubulin levels on both sides of the KT (*p* = 0.016; *p* = 0.025) (Figure 4K, green circles) relative to controls (Figure 4K, black circles). This is also reflected in the tubulin ratio (M/P) (Figure S3A). Representative images of cells overexpressing MCAK/Kif2C are shown in Figure S3D.

In the case of MCAK/Kif2C-depleted cells, we hypothesize that impaired congression, increased IKD, and lower K-fiber turnover reflect more lateral interactions with merotelic bundles of MTs present near the M faces of centromeres, as defined by Hec1 label. In contrast, excess MCAK/Kif2C appears to lower tubulin levels on both faces of the centromere yet still appears to decrease K-fiber turnover as indicated by increased acetylation. We believe this stems from increased IKD as sisters cannot coordinate with each other because K-fiber MTs are unable to polymerize in response to higher tension^22^ due to higher MCAK/Kif2C levels at metaphase. These models are diagrammed in Figure 4L.

## DISCUSSION

Both MCAK/Kif2C loss and OE produce identical downstream phenotypes. Both treatments increase IKD, which is responsible for increased stabilization of KT fiber MTs in both conditions. Excess stabilization of k-fibers has the potential to promote the anaphase segregation errors seen in both of these conditions. We also show that the increased IKD we see in both treatments has different mechanistic origins and distinct tubulin ratios on each side of the centromere. We do not yet have the spatial resolution to investigate this in live cells. For example, we hypothesize that surplus centromere-associated MCAK/Kif2C leads to high IKD due to excess depolymerization of the KT-bound MTs. It has been shown that MCAK/Kif2C is primarily responsible for promoting end-on MT interactions.^33^ Furthermore, MT end-bound MCAK/Kif2C can use depolymerization activity to do work and exert force on centromeres.^34^ Importantly, MT growth in KT-attached MTs is promoted by high tension at the KT.^22^ In cells with excess levels of MCAK/Kif2C on the centromere, this effect would be prevented, leading to a lack of coordination between sister centromeres. We note that levels of centromeric MCAK/Kif2C naturally decrease from prometaphase to metaphase (Figure S3B). The reason for this is unknown, but it may be to allow for coordinated oscillatory movements in chromosomes with multi-MT KTs.^22^

In contrast, when MCAK/Kif2C is completely missing prior to chromosome alignment, we hypothesize that there is a poor force balance between sister-centromeres due to increased numbers of lateral MT interactions. This leads to increased uncoordinated movement and increased IKD as sisters ‘‘fight’’ each other. Unlike the case with excess MCAK/Kif2C, there is less translocation, so the lack of coordination is less obvious. Live tracking of MCAK/Kif2C-depleted centromeres shows decreased oscillatory co-ordination between sisters (Figure S3C). At this time, the simplest interpretation is that loss of MCAK/ Kif2C promotes excess lateral KT-MT interactions whereas excess MCAK/Kif2C suppresses polymerization of KT-bound MTs. In both cases, these conditions disrupt KT coordination, leading to excess tension and decreased K-fiber turnover.

## Limitations of the study

There are two key limitations to this study. One limitation of the study is that super high-resolution microscopy or electron microscopy of a large number of KTs is required to make a final judgment call on whether lateral MT interactions are restricted to low MCAK/Kif2C conditions. This would support our hypothetical distinction between KTs with excess versus abnormally low levels of MCAK/Kif2C. A final limitation is that we were unable to directly measure k-fiber turnover in conditions of high versus low levels of MCAK/Kif2C. We expect both conditions to exhibit low turnover relative to control cells, which would support our indirect measurements of k-fiber acetylation.

## RESOURCE AVAILABILITY

### Lead contact

- Requests for further information and resources should be directed to the lead contact, Linda Wordeman.

### Materials availability

- Microscopy data reported in this paper will be shared by the lead contact upon request.

### Data and code availability

- Microscopy data reported in this paper will be shared by the lead contact upon request.
- This study produced no code.
- Any additional information may be requested from the lead contact.

## ACKNOWLEDGMENTS

The authors thank the Seattle Mitosis Club and Chip Asbury for feedback on this study. We also thank the UW Pathology Flow Cytometry Core Facility for cell sorting assistance and Nikon for live imaging assistance and advice.

This study was supported by the NIH National Institute of General Medicine R01 GM145567 (L.W.) and UW Royalty Research Fund (J-J. R).

## AUTHOR CONTRIBUTIONS

Conceptualization, L.W. and M.W.; methodology, L.W., J.J.V., and M.W.; investigation, L.W. and M.W.; writing – original draft, L.W.; writing – review and editing, L.W., M.W., and J.J.V.; funding acquisition, L.W.; resources, L.W. and M.W.; supervision, L.W.

## STAR METHODS

Detailed methods are provided in the online version of this paper and include the following:

- KEY RESOURCES TABLE
- EXPERIMENTAL MODEL AND STUDY PARTICIPANT DETAILS
- METHOD DETAILS
  - Cell lines and plasmids
  - Immunofluorescence
  - Live imaging
  - Image analysis and statistics
- QUANTIFICATION AND STATISTICAL ANALYSIS
- ADDITIONAL RESOURCES

## SUPPLEMENTAL INFORMATION

Supplemental information can be found online at https://doi.org/10.1016/j.isci.2026.116122.

Received: November 13, 2025

Revised: April 14, 2026

Accepted: May 11, 2026

## STAR METHODS

### KEY RESOURCES TABLE

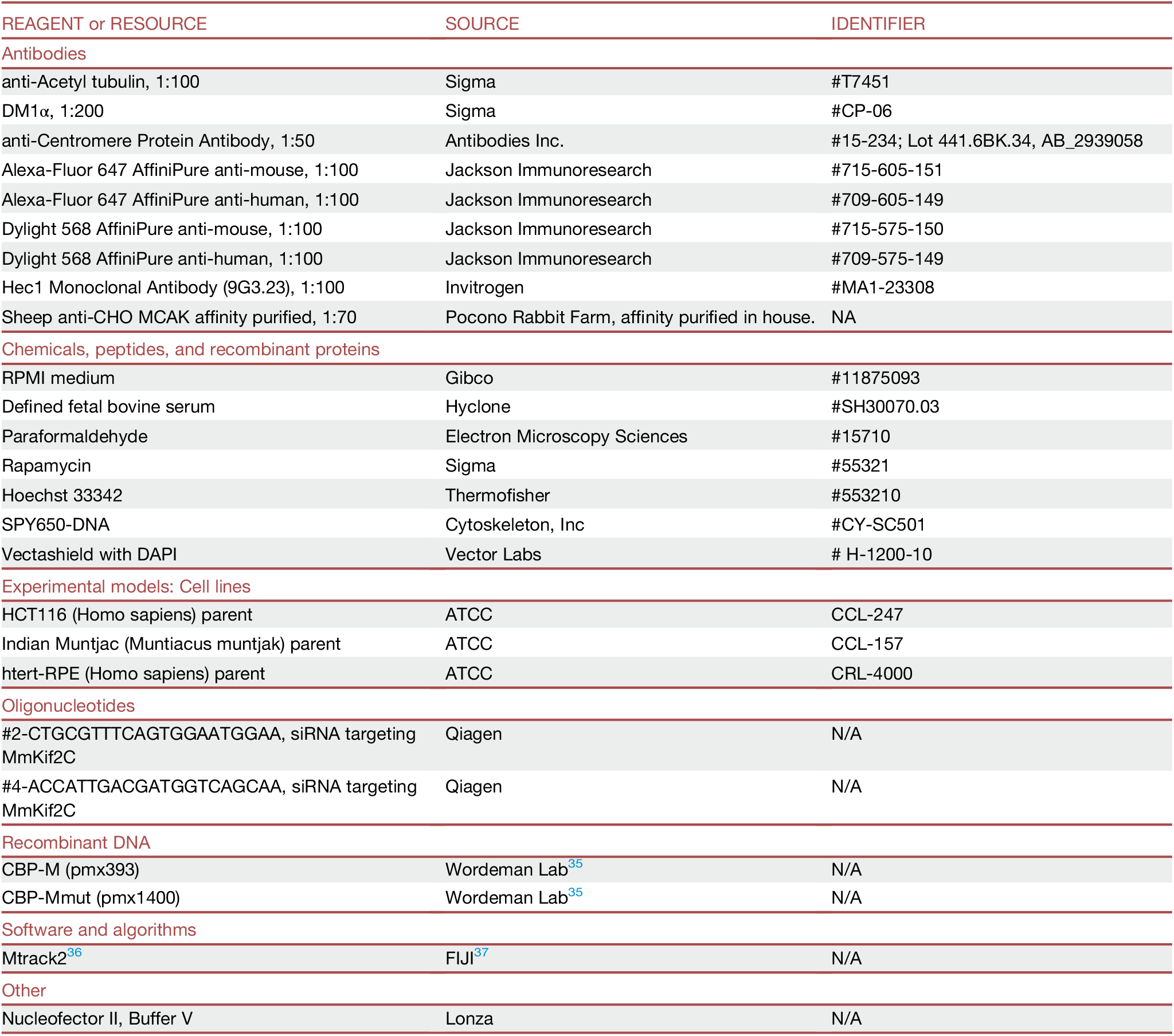

### EXPERIMENTAL MODEL AND STUDY PARTICIPANT DETAILS

Please list here under separate headings all the experimental models/study participants (animals, human participants, plants, microbe strains, cell lines, primary cell cultures) used in the study. For each model, provide information related to their species/strain, genotype, age/developmental stage, sex (and gender, ancestry, race, and ethnicity if reported for human studies), maintenance, and care, including institutional permission and oversight information for the experimental animal/human participant study. The influence (or association) of sex, gender, or both on the results of the study must be reported. In cases where it cannot, authors should discuss this as a limitation to their research’s generalizability.

## METHOD DETAILS

### Cell lines and plasmids

CRISPR engineered HCT116 and RPE-hTERT cells were cultured at 37^°^C and 5% CO_2_ in RPMI medium (#11875093; Gibco) with 2 mM L-Glutamine supplemented with 10% defined fetal bovine serum (#SH30070.03; Hyclone). Indian Muntjac cells were obtained from ATCC (CCL-157) were cultured at 37^°^C and 5% CO_2_ in RPMI medium (#11875093; Gibco) with 2 mM L-Glutamine supplemented with 20% defined fetal bovine serum (#SH30070.03; Hyclone). The cherry-MCAK/Kif2C, CBP-M and CBP-Mmut constructs are described here.^35^ Transfections were performed using a Nucleofector II (Lonza) according to the manufacturer’s protocol. Detailed instructions on the preparation of CRISPR cell lines are provided here.^11^ Cells were plated onto 35-mm glass-bottom dishes (MatTek) for live imaging or 12 mm, 1.5 coverslips for immunofluorescence (#72290-04, Electron Microscopy Sciences).

### Immunofluorescence

For spinning disk confocal imaging, cells were fixed in prewarmed 4% PFA (#15710; Electron Microscopy Sciences) in PBS for 10 min at 37^°^C. Cells were then incubated overnight at 4^°^C with anti-Acetyl tubulin (#T7451; Sigma), DM1α (#CP-06; Sigma) or anti-ACA antibodies (#15-234; Antibodies Inc.). Antibody binding was visualized by incubation of the coverslips in Dylight 568 secondary antibodies (Jackson) at room temperature. Coverslips were then washed three times in PBS and mounted on slides in Vectashield (VectorLabs).

### Live imaging

Live imaging was performed on a Nikon CREST spinning disk confocal microscope with 37^°^C heat block stage (Dagan Corp.). Three z-planes were imaged at 2 or 5 min. intervals. Five separate imaging sessions of full fields containing 5 to 10 individual mitotic cells were tracked. Laser power was ≤5%. Alternatively, some live cell experiments were imaged in a Nikon Biostation IM-Q (Nikon). Cells were treated with 1µm rapamycin (#553210; Sigma) for 4 hours prior to imaging. Media was washed out three times and replaced with fresh 37^°^C media with no drugs. Relocalization is not reversible so no drugs are necessary while imaging. Imaging was performed for 2 hours using 5 min. timepoints. For tracking of natural congression and mitotic timing, cells were imaged for 24 hours at 5 min. timepoints. Briefly, cells were timed from nuclear envelope breakdown, which was visible in the DNA channel, to the first establishment of a metaphase plate of at least three microns in width. This was scored as time in congression (NEB to metaphase). A few cells would go on to lose this plate leading to an increase in time scored as metaphase timing. This was relatively rare and the majority of cells were measurably consistent in metaphase-to-anaphase timing. Microtubule assembly assays were performed as described.^19,27^ Live sister-kinetochore tracking was performed as described^38^ using TrackMate or MTrack2 as needed (see below).

### Image analysis and statistics

All images were processed using Nikon NIS-Elements (Nikon) or FIJI software (https://fiji.sc/). Sister centromere movements were tracked by hand using MTrack2 (FIJI). Individual centromeres (meaning not sisters) were tracked using TrackMate (FIJI;^36^). Congression was tracked by hand, frame-by-frame from nuclear envelope breakdown to anaphase and the presence or absence of lagging chromosomes during anaphase was recorded (labeled with Hoechst 33342 (#H1399; ThermoFisher) or SPY650-DNA (#CY-SC501; Cytoskeleton, Inc.). Metaphase was timed as the onset of alignment. When lagging chromosomes were present it was not feasible to count them so the data is scored +/- for the presence of lagging chromosomes. Integrated density of fluorescence was measured in the acetyl-tubulin channel using a circle of consistent size (for all cells measured) that is 0.2µm from the polar end of the spindle and background subtracted from cytoplasmic non-spindle fluorescence. In the Indian muntjac cells the Feret’s diameter and tubulin ratio was measured for the large X centromere only and each measurement corresponds to one sister centromere in one cell. The centromere was traced by hand and the shortest distance between the beginning and end point was measured as Feret’s diameter. The Feret diameter (sometimes called the caliper diameter) is the measure of an object’s size along a specified direction. The centromere fluorescence is traced as a line with the Feret diameter being the shortest distance between the beginning and end of the line. In this application, the maximum Feret diameter of a properly aligned X-chromosome centromere is 1.0. This corresponds to a straight line oriented perpendicular to the spindle axis extending from each visible edge of the centromere. Measurements that are <1.0 correspond to increasing levels of deformation of the centromere from a straight plate oriented perpendicular to the long spindle axis to a stretched or merotelic attachment oriented longitudinal to the long spindle axis. The tubulin ratio was measured from fixed Indian muntjac cells using a SUMMED Z-stack of 3 sections of 0.5 microns. tubulin integrated density was measured for P and M using identical sized boxes placed on either side of the centromere relative to the longitudinal spindle axis.

Fluorescence intensity is measured as arbitrary integrated density units. Measurements within the spindle were background subtracted from cytoplasmic fluorescence. For individual centromeres fluorescence intensity was measured as the brightest point on the centromere and also background subtracted for cytoplasm fluorescence. Acetyl-tubulin and tubulin were quantified as the integrated density of a consistently sized circle of 2µm by 2µm located 0.2 µm from the spindle pole in a summed projection of 21 Z-planes. IKDs were measured by hand using Fiji from stacks consisting of 21 Z planes on only those centromeres that could clearly be identified as sister-centromeres.

### QUANTIFICATION AND STATISTICAL ANALYSIS

For all statistics, the data distributions were assumed to be non-normal and subjected to Mann-Whitney t-tests (Prism 10; Graphpad). Graphs are presented with medians and 95% confidence intervals shown.

## ADDITIONAL RESOURCES

Image J/FIJI software https://fiji.sc/.

