## Supplemental Figures for "MCAK/Kif2C centromeric activity level tunes K-fiber turnover through distinct pathways"

**iScience, Volume ■ ■**

### Supplementary Figure S1

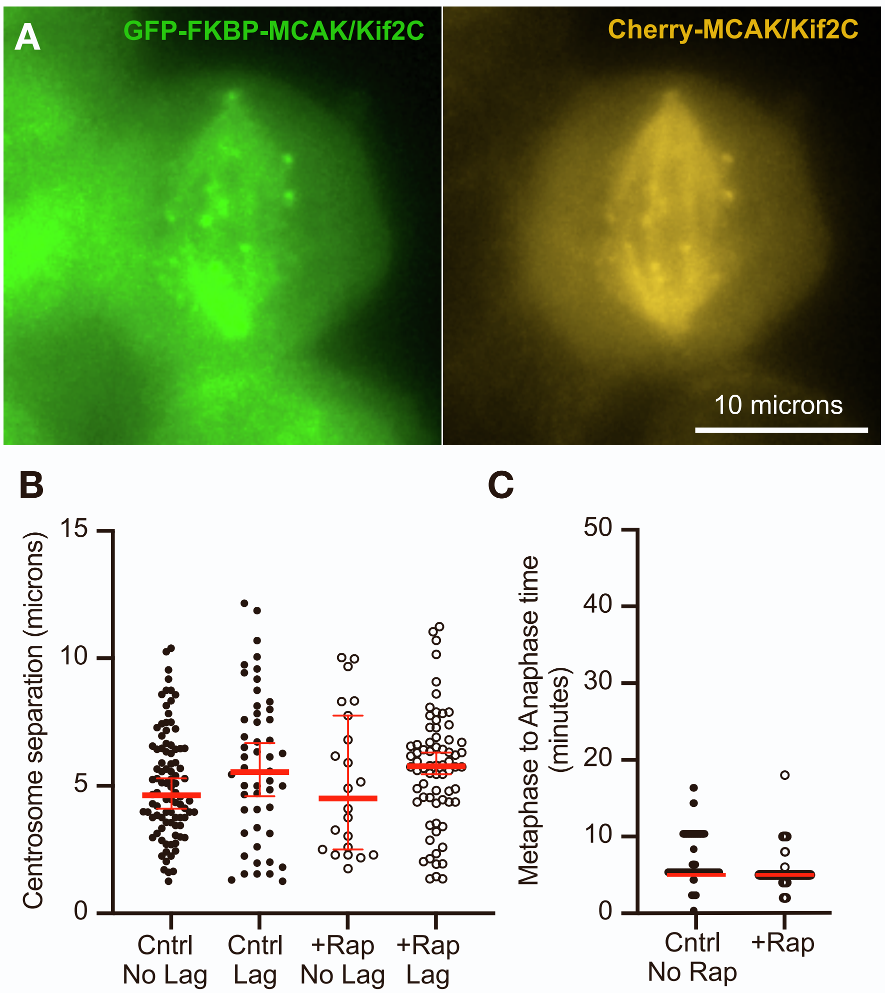


Figure S1. Addition of excess cherry-MCAK/Kif2C has no effect on endogenous GFP-FKBP-MCAK/Kif2C. A. Endogenous GFP-FKBP-MCAK/Kif2C in CRISPR-engineered cell (left) that has been transfected with additional Cherry-MCAK/Kif2C (right). Additionally, Loss of MCAK/Kif2C had no effect on extent of centrosome separation or metaphase to anaphase timing. B. There is no significant difference in centromere separation prior to NEB in cells that manifest lagging chromosomes versus those that do not (black circles). Furthermore, addition of rapamycin to relocalize MCAK/Kif2C does not alter the extent of centrosome separation (open circles). C. Timing of the progression from metaphase to anaphase was not significantly different in cells exposed to rapamycin (open circles) to relocalize MCAK/Kif2C as compared to cells with MCAK/Kif2C present (black circles). This proved true for either naturally congressing cells (left) or cells recovering from monastrol treatment (right).

### Supplementary Figure S2

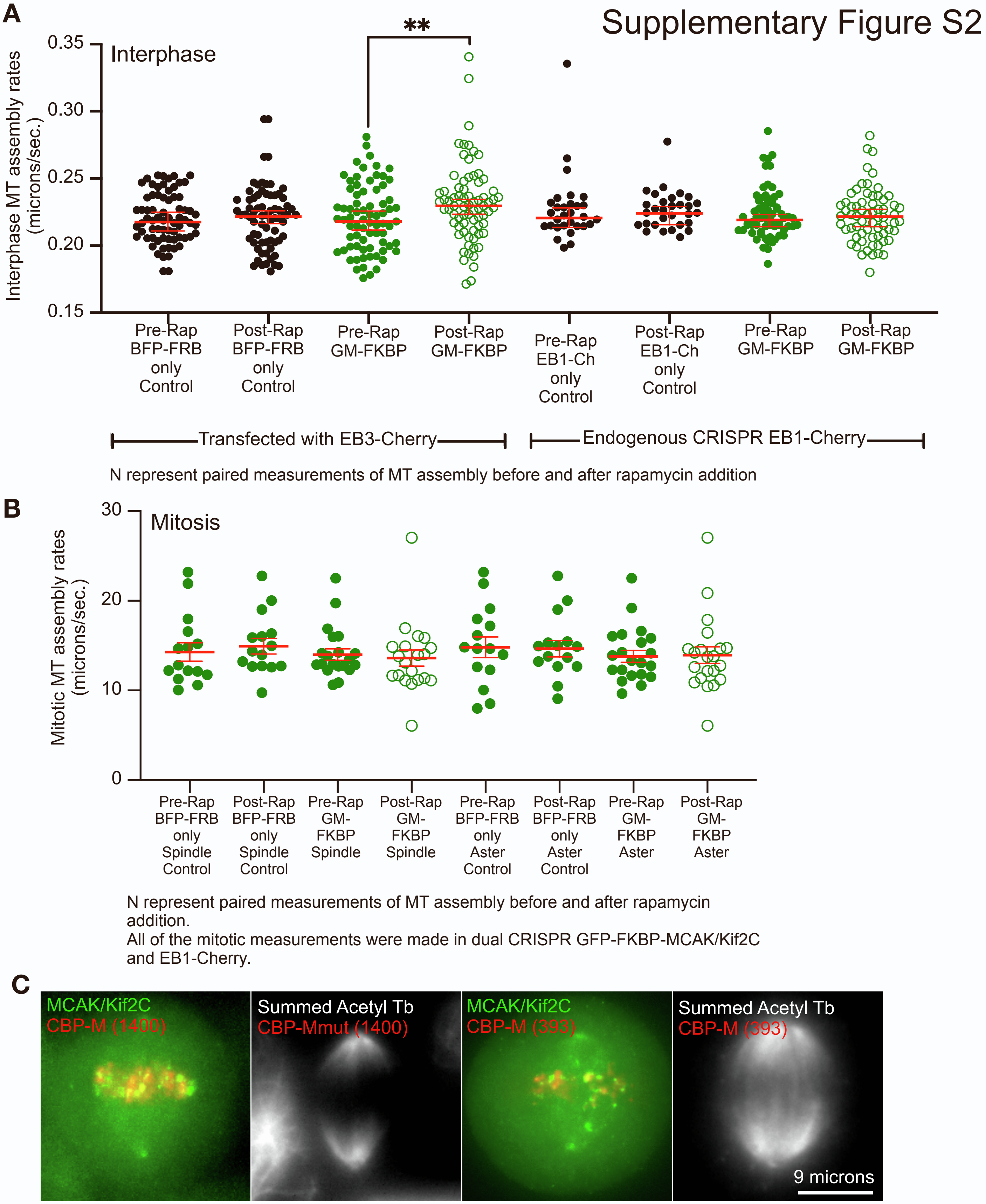


Figure S2. Relocation of MCAK/Kif2C does not affect MT assembly rates unless you measure rates with EB3. A. Paired rate measurements of interphase cells transfected with EB3-cherry. Controls (black circles) are cells with no GFP-FKBP-MCAK/Kif2C. Cells with relocalized MCAK/KIf2C (open green circles) do not differ in MT assembly rates from pre-rapamycin (green circles) cells unless EB3-cherry is present (p=0.0010). B. Paired rate measurements of MT assembly in dual CRISPR GFP-FKBP-MCAK/Kif2C, EB1-cherry cells either within the mitotic spindle (between the centrosomes) or in the asters. Relocalization of MCAK/KIf2C has no effect on MT assembly rates in the spindle or asters (compare pre-treated, green circles, cells with cells that have relocalized in rapamycin, open green circles). C. CBP-M constructs do not disrupt GFP-FKBP-MCAK/Kif2C distrubution in CRISPR cells. They localize to mostly different areas.

### Supplementary Figure S3

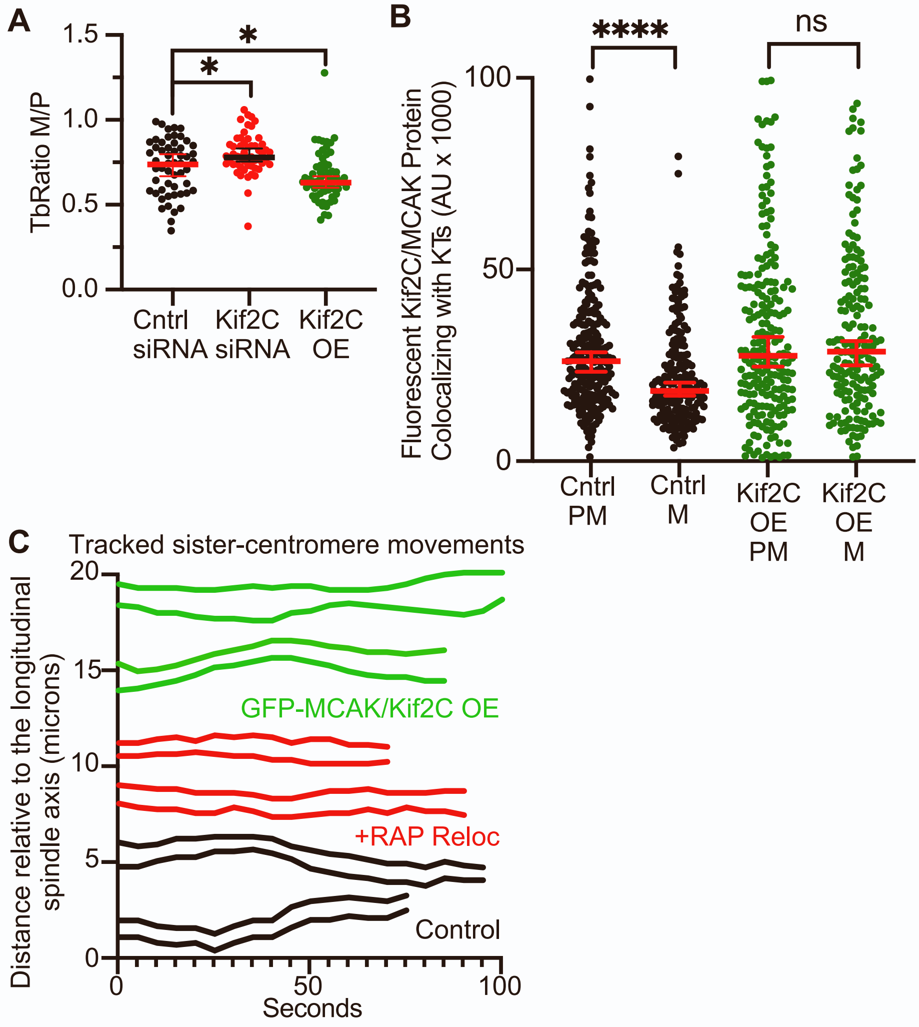


Figure S3. Tubulin ratio has the potential to affect centromere motility. A. Tubulin ratio (M/P) for the large X centromere is higher in MCAK/Kif2C depleted cells (* p=0.0243) and lower in MCAK/Kif2C OE cells (* p=0.0153). B. Centromere-associated MCAK/Kif2C becomes naturally significantly (**** p<0.0001) lower as cells progress from prometaphase to metaphase in Control cells (black circles). There is no significant difference (ns p=0.564) between centromere-associated MCAK/Kif2C levels in cells the over-express MCAK/Kif2C (green circles). C. Individually tracked sister centromere pairs in live cells. Centromeres in cells over-expressing MCAK/Kif2C (green) appear to lose co-ordination at times leading to large IKD but they still oscillate. Centromeres depleted of MCAK/Kif2C (red) appear uncoordinated and do not oscillate well. Control cells (black circles) exhibit typical oscillations with good co-ordination between sister centromeres. Separation between the traces corresponds to the measured IKD. All comparisons are Mann-Whitney t-tests
